# Differences in DNA methylation of white blood cell types at birth and in adulthood reflect postnatal immune maturation and influence accuracy of cell type prediction

**DOI:** 10.1101/399279

**Authors:** Meaghan J Jones, Louie Dinh, Hamid Reza Razzaghian, Olivia de Goede, Julia L MacIsaac, Alexander M. Morin, Kristina Gervin, Raymond Ng, Liesbeth Duijts, Menno C van Zelm, Henriëtte A Moll, Robert Lyle, Wendy P Robinson, Devin C Koestler, Janine F Felix, Pascal M Lavoie, Sara Mostafavi, Michael S Kobor

## Abstract

**Background:** DNA methylation profiling of peripheral blood leukocytes has many research applications, and characterizing the changes in DNA methylation of specific white blood cell types between newborn and adult could add insight into the maturation of the immune system. As a consequence of developmental changes, DNA methylation profiles derived from adult white blood cells are poor references for prediction of cord blood cell types from DNA methylation data. We thus examined cell-type specific differences in DNA methylation in leukocyte subsets between cord and adult blood, and assessed the impact of these differences on prediction of cell types in cord blood.

**Results:** Though all cell types showed differences between cord and adult blood, some specific patterns stood out that reflected how the immune system changes after birth. In cord blood, lymphoid cells showed less variability than in adult, potentially demonstrating their naïve status. In fact, cord CD4 and CD8 T cells were so similar that genetic effects on DNA methylation were greater than cell type effects in our analysis, and CD8 T cell frequencies remained difficult to predict, even after optimizing the library used for cord blood composition estimation. Myeloid cells showed fewer changes between cord and adult and also less variability, with monocytes showing the fewest sites of DNA methylation change between cord and adult. Finally, including nucleated red blood cells in the reference library was necessary for accurate cell type predictions in cord blood.

**Conclusion:** Changes in DNA methylation with age were highly cell type specific, and those differences paralleled what is known about the maturation of the postnatal immune system.

## Background

One of the main established roles for DNA methylation (DNAm) is in development, where it contributes to the functional maturation, lineage commitment and fate of cells.^1^ This has two important implications; the first is that DNAm within a given cell type will change over time as cells differentiate and function develops.^2^ The second is that different types of terminally differentiated cells will have very distinct DNAm profiles.^3,4^ As the age of individuals and cell type are two of the major determinants of DNAm variability, analyses of DNAm data must carefully consider those variables.^5,6^ Due to an important role of DNAm in development, close assessment of developmental processes, by identifying specific genes or genomic regions that change with age.

This is of particular interest in blood, where development of the immune system in early life is linked to long term health outcomes, and so the analysis of the changes in DNAm from birth to adulthood may provide insights into how the immune system matures. Umbilical cord blood is an important and much utilized research tissue, as it is easy to collect from the umbilical cord post-delivery, and thus many studies have assessed DNAm in relatively large numbers of cord blood samples.^7,8^ Cord blood is very distinct from adult blood, as it contains a much greater abundance of nucleated red blood cells (nRBC) expressing unique proteins such as fetal hemoglobin, as well as functionally distinct myeloid and lymphoid cells^9,10^ These distinct functions reflect the greater reliance on innate immunity in newborns, as adaptive immune cells requires exposure to pathogens in order to mature and generate functional memory^11,12^ Thus, one might expect that innate immune cells such as granulocytes, monocytes, and NK cells, would be more similar over development than adaptive immune cells like B and T cells. However, this relationship is more complex, with differences observed even in the function of innate immunity between newborns and adults, indicating that the functionality of specific innate cell types also changes over development^13,12,14^.

These biologically meaningful differences in function are likely to be reflected in DNAm changes over developmental time, and thus can cause complications for the analysis of DNAm data, as computational tools designed for use in adult blood may not function as well for blood from children or newborns. An example of this is cell-type deconvolution, which is one of the major tools used to account for inter-individual differences in cell type composition in mixed tissue samples, such as blood, when more direct measures are not available.^15-19^ Failing to account for these inter-individual differences in cell type composition can lead to both false positive and false negative results in epigenetic association studies, and therefore accurate implementation of this tool in a developmental context is essential.^2,5^ Perhaps not surprisingly, as the most commonly used tool was designed for adult references, it performs poorly on cord blood data.^20-24^ In an attempt to address this problem, three different reference datasets for cord blood to create developmental stage specific libraries have been published, but validation studies using these updated references only partially close the gap between adult and cord blood prediction accuracy.^20,22,25^

In this study, we compared DNAm profiles of purified leukocyte subsets from cord and adult blood, with the goal of further understanding the biological differences in each cell type as they mature. Using these insights, we then tested specific assumptions of existing deconvolution methods for estimating cell type proportions in cord blood, modified the algorithm to account for the differences between cord and adult, and evaluated the prediction accuracy on two data sets. We showed that differences between cord and adult blood cell types reflected the functional maturation of the immune system, and these differences must be incorporated into the design of methods to be used on DNAm data.

## Results

### Cell type-specific DNA methylation in adult and cord blood

Previous reports have shown that adult references poorly predict cell types in cord blood.^20-22^ We hypothesized that differences in DNAm between cord and adult blood might impact the performance of cell type deconvolution, and so compared patterns of DNAm in the adult and cord blood reference data sets.^3,20,25^ In order to take advantage of as many samples as possible, we combined two previously published cord blood data sets, and compared them to a publicly available adult blood data set (Table 1).^3,20,25^ All three data sets were generated from Fluorescence Activated Cell Sorting (FACS)- isolated white blood cell types from healthy donors, resulting in 6 sets of adult blood and 18 sets of matched 450k cord blood DNAm profiles. All three sets were combined and processed together, and after processing and filtering, 428,688 probes remained. Visualization by hierarchical clustering of all CpGs analyzed showed that samples grouped first by myeloid (granulocytes, monocytes) versus lymphoid (B, T, NK cells) lineage, then by age, and finally by specific cell type (Figure 1A). Adult lymphocytes were the most distinct group, followed by nRBCs. In cord blood samples, CD4 and CD8 T cells clustered in one large group, paired by individual as opposed to cell type. This indicated that the influence of genotypic variation within our study population outweighed the influence of cell type on DNAm patterns of CD4 and CD8 T cells in cord blood. To further test this, we performed a silhouette analysis, with cell types as clusters. Consistent with expectations, all cell types clustered relatively well, with the exception of CD8 T cells, where cord CD8 T cells did not cluster well with adult CD8 T cells (Figure S1).

**Table 1:** Summary of data sets used in this study

|  | Reinius | Gervin | de Goede |
| --- | --- | --- | --- |
| Age | Adult | Cord | Cord |
| % female | 0 | 54.5 | 71.4 |

| N |  |  |  |
| --- | --- | --- | --- |
| CD8T | 6 | 11 | 6 |
| CD4T | 6 | 11 | 7 |
| B | 6 | 11 | 7 |
| NK | 6 | 11 | 7 |
| Granulocyte | 18 | 11 | 7 |
| Monocyte | 6 | 11 | 7 |
| nRBC | 0 | 0 | 7 |

**Figure 1:**
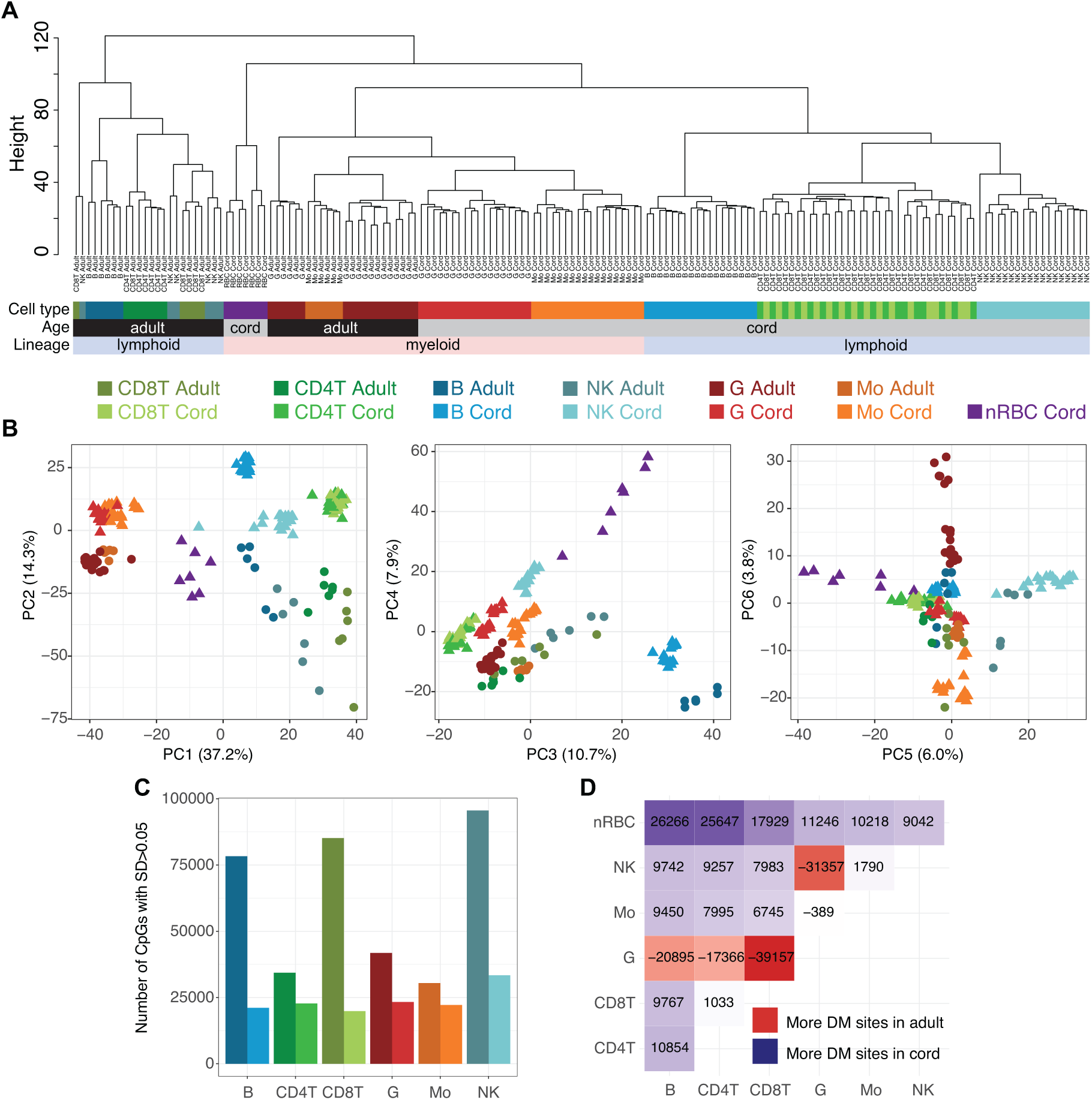
DNA methylation patterns of cord blood cell types were highly distinct from the corresponding cell types in adult blood. **A)** Dendrogram of 162 samples, n=5 for each adult cell type, n=18 for cord B cells, CD4 T cells, granulocytes, monocytes, and NK cells, n=17 for CD8 T cells, and n=7 for nRBCs. Samples clustered first by lineage (pink = myeloid, pale blue= lymphoid), then by age (black = adult, grey = cord), and then by specific cell type (colour scale below). **B)** The first six principal components of the data set in A, where circles are adult samples and triangles are cord blood samples, colours as above, and percent of variance indicated on the relevant axis. **C)** Number of sites in each cell type with an SD>0.05 in adult and cord cell types. See full counts of variable sites for all cell types and cell mixtures in Table S1. **D)** Heatmap showing number of sites that distinguish between each pair of cell types in adult versus cord data (adult nRBC values were set to zero). The red colour indicates that more sites distinguish these cell types in adult and purple indicates that more sites distinguish these cell types in cord.

Next, we used Principal Component Analysis (PCA) to determine how the patterns of variability in DNAm differed between cord and adult blood cell types. We first examined the first six principal components (PCs), accounting for more than 80% of the variance and which separate the different cell types. The first PC, accounting for 37% of the variance, separated the myeloid and lymphoid lineages (t-test p<1×10^−16^), with distinct clustering of cord and adult samples in the second PC (t-test p<1×10^−16^), accounting for 14% of the variance (Figure 1B). Myeloid cell types clustered more closely than lymphoid cell types across PCs, perhaps reflecting a relative functional and lineage proximity. These findings were consistent with the results of our hierarchical clustering analysis. Next, we visually examined the spread of PC scores within a cell type, an indication of how similar the samples within a cell type are to one another. Across both of the top PCs, adult lymphoid cell types showed greater variability compared to myeloid cell types (Figure 1B). The variability within adult lymphoid cells was also higher than their corresponding cord blood cell types in these first PCs, which may reflect an increasing proportion of differentiated effector and memory T and B cells due to antigen exposure over lifespan.

To quantify the observed differences in variability between cord and adult blood observed by PCA, we examined the number of variable sites within each cell type. We hypothesized that a high number of variable sites in any particular cell type might make it more difficult to identify tissue-specific sites, as variable sites within a cell type are unlikely to be good cell type markers. Due to different sample numbers, we compared the adult samples only to one sorted cord blood data set (de Goede), which have similar sample sizes (n=6 adult, n=7 cord), but includes nRBCs. We defined a variable probe as having a beta value standard deviation greater than 0.05 (Table S1). Notably, nRBCs exhibited a large number of variable probes (77,888). All cell types showed more variable sites in adult than in cord, and B cells, CD8 T cells, and NK cells showed considerably more than CD4 T cells, monocytes and granulocytes. While overall, the total numbers of variable sites were likely not high enough to influence the accuracy of cell type prediction, higher variability in adult lymphoid cell types might be reflective of inter-individual differences in adaptive immunity (Figure 1C, raw counts in Table S1).

We next determined whether cell type pairs were more or less difficult to distinguish from one another in cord blood as compared to adult blood. To do this, we extracted the number of differentially methylated probes within cord or adult whole blood using pairwise comparisons for all cell types with a stringent nominal p value of 1×10^−7^. As expected, we found different candidate cell type DNAm markers between the two ages, but also noted important differences in the numbers of CpGs that can distinguish between cord and adult blood cell types (Figure 1D). All pairs, except those involving granulocytes, had more sites distinguishing pairs in cord than in adult samples. This may be because cord samples were less variable than adult within a cell type as observed in both PCA and the number of variable sites, making it easier to distinguish one cell type from another.

To identify cell type specific DNAm differences between cord and adult blood, we performed an epigenome-wide association study (EWAS) between cord and adult in the six common cell types (CD4 T, CD8 T, NK, B, granulocyte and monocyte) and the two cell mixtures (whole blood and mononuclear cells). As expected, a large number of CpGs were differentially methylated between cord and adult blood at a p value of 1×10^−7^ and a mean DNAm difference of 10% (min 2989 for monocytes, max 9885 for granulocytes, Figure 2A, Figure S2, Table S2). In the purified cell types, 588 CpGs were differentially methylated between cord and adult samples in all six types, and 397 and 2062 CpGs were lymphoid or myeloid specific, respectively (Figure 2A, full overlap in Table S3).

**Figure 2.**
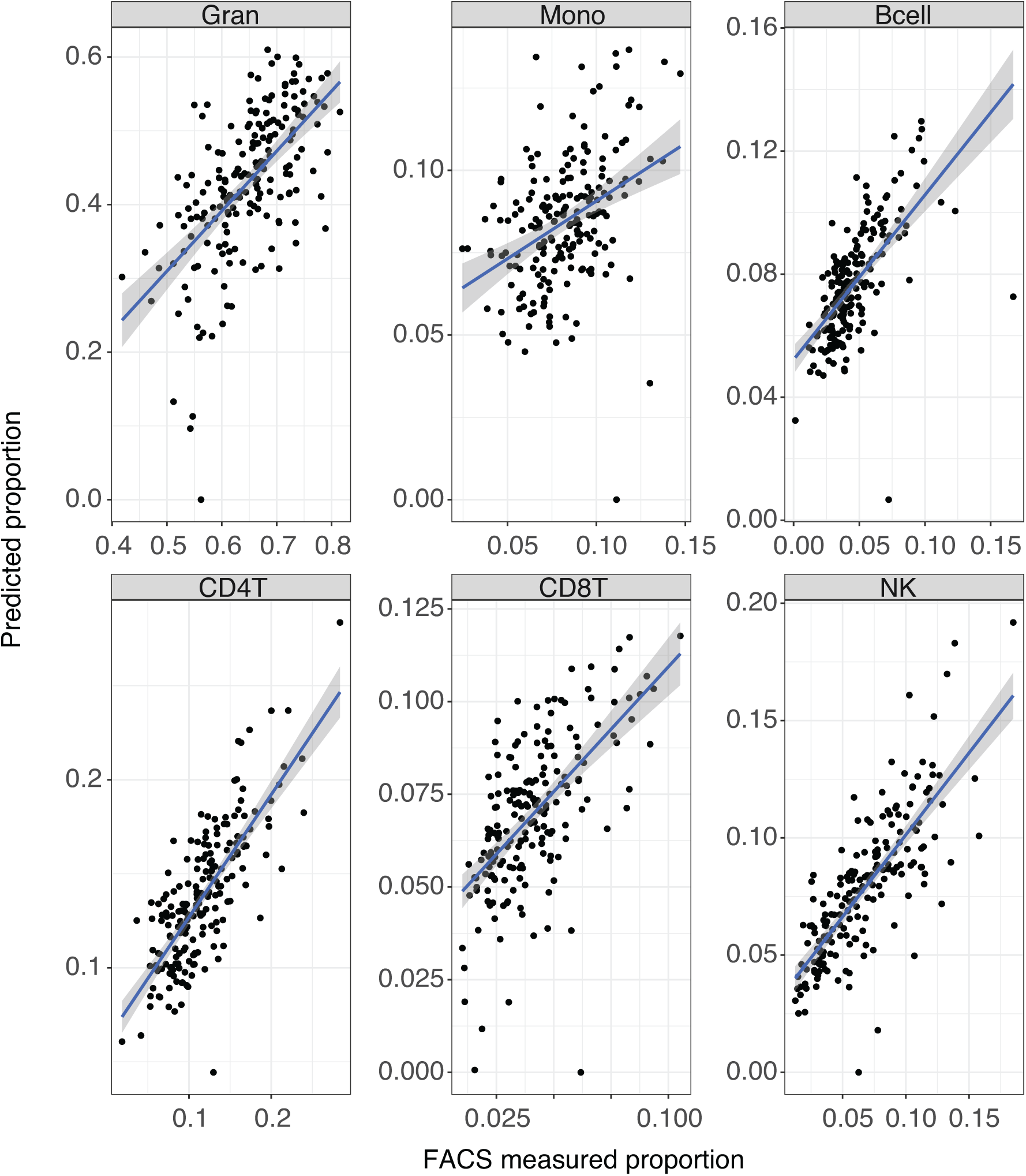
DNA methylation differences between cord and adult blood cells by cell type and genomic location. We performed an EWAS comparing cord to adult samples for each cell type, retaining sites with a p value <1×10^−7^ and a mean DNA methylation difference >10% (visualized in Figure S2). **A)** The number of significantly differentially methylated CpGs between cord and adult blood in the six cell types. Significant CpGs in all cell types are in grey (N=588), lymphoid specific (N=397) or myeloid specific (N=2062) CpGs are in pale blue or pink, respectively, and the remaining CpGs are in the colour of that cell type. Note that these might not all be unique to that cell type, but are neither common, nor specific to lymphoid or myeloid cells. Total pairwise overlap numbers are in Table S2. **B)** Number of CpGs in mixed tissues which were differentially methylated (N=2558 and N=1993 in whole blood and mononuclear cells, respectively), and overlap with the differentially methylated sites in each cell type. The number of sites common across all cell types and cell mixtures (N=507 out of 588 in grey in part A) was subtracted from the total number. **C)** Proportion of differentially methylated CpGs in each cell type and cell mixture that fall within each of the four CpG island classes (HC = high density, IC = intermediate density, ICshore = CpG island shore LC=low density). **D)** Proportion of differentially methylated CpGs in each cell type and cell mixture that fall within five common genomic features. Sites not annotated to a specific region are not shown.

In an effort to determine how much of these differences were due to genetic effects, given that our cord and adult samples were not from the same individuals, we examined the overlap with CpGs previously identified as being associated with genotype (mQTLs) in cord blood in the ARIES data set (Table S3).^8^ We hypothesized that genetic effects would be observed at highest proportion in those sites which were differentially methylated between cord and adult samples in all cell types, as cross-tissue genetic effects seem to be more frequent than tissue-specific genetic effects.^26,27^ The results from these analyses revealed that between 9 and 11% of the myeloid- and lymphoid-specific differentially methylated sites were currently reported mQTLs. In the individual lymphoid cell types, 6-12% of the differentially methylated CpGs between cord and adult blood were cord blood mQTLs. Interestingly, the myeloid cell types showed a very different pattern, where 18% of the differentially methylated CpGs in granulocytes and 79% in monocytes were cord blood mQTLs. This result was surprising, as it implies that many of the differentially methylated CpGs between cord and adult blood in monocytes might be mQTLs, despite the fact that it already had the smallest number of differentially methylated sites among cell types. This could have interesting implications for future assessment of genetic influences on cell type specific DNAm.

Next, we examined the differentially methylated sites between cord and adult in two commonly used cell mixtures; whole blood, which contains all cell types and granulocytes are by far the most prevalent and blood enriched for mononuclear cells, which primarily removes granulocytes, leaving CD4T cells as the most prevalent cell type. We hypothesized that the differences between cord and adult in these mixtures would be influenced by the underlying cell proportions, meaning that differentially methylated CpGs in each mixture would overlap most with the most prevalent cell type in that mixture. This was indeed observed, with differentially methylated sites in mononuclear cells overlapping most with those sites which were differentially methylated in CD4 T cells and, and likewise whole blood sites overlapped most with granulocyte sites (Figure 2B).

Finally, we examined the genomic feature locations of the differentially methylated sites between cord and adult in all cell types by mapping each CpG to CpG island class (HC: high density CpG island, LC: low density CpG island, IC: intermediate density CpG island) and genomic features. In NK and B cells, more CpGs mapped to high density (HC) islands and less for low density (LC) islands. CD4 T cells showed the opposite pattern. Overall the cell types were quite consistent in enrichment for CpG island status (Figure 2C). Few differences between the cell types and mixtures were observed for enrichment of the six genomic regions (1^st^ exon, 3’UTR, 5’UTR, gene body, TSS200, and TSS1500) investigated (Figure 2D).

### Probe type selection method and inclusion of nRBCs influenced cell type prediction accuracy in cord blood

Given that DNAm showed substantial differences between cord and adult blood in both variability and at specific sites across the genome, we next identified which parts of the deconvolution algorithm might be affecting the accuracy of predictions in cord blood. To do this, we used a validation data set of 24 whole cord blood samples from which both DNAm measurements and matched cell counts determined by flow cytometry were available. First, we applied the existing deconvolution algorithm from the minfi package using default settings to the test data using the adult references, and repeated the same prediction using the cord references that included nRBCs. The results from these analyses revealed moderate prediction of cord blood cell types using adult references, and slightly improved predictions using the de Goede cord blood references, in agreement with previous studies (Figure S3).^20-22^

We then hypothesized that the method for selecting sites to use in deconvolution could influence its prediction accuracy, as the observed differences between cord and adult DNAm mean that the method created for adult blood may be less effective on cord blood. Several selection heuristics have been proposed and modified over the past few years.^28,29^ The original method selected the top 50 probes that display higher and lower DNAm in each cell type according to their effect size, for a total of 100 probes per cell type. In cord blood, we replicated a previous finding that for monocytes in particular, this selection method chooses many sites that do not distinguish between monocytes and other cell types, as there are less than 20 monocyte markers that have higher DNAm in monocytes than other cell types (Figure S4).^22^ This means that in cord blood, forcing probe selection to include an equal number of higher and lower methylated probes would adversely affect monocyte prediction at the very least, which would in turn reduce the accuracy of prediction for the other cell types.

Next we examined the predictions of nRBCs, which can account for up to 25% of the nucleated cell composition in cord blood, and possibly more of the DNAm signal due to cell free DNA from nRBCs that had already extruded their nuclei.^25^ As previously reported, nRBCs have a unique DNAm profile in cord blood, quite different from the typical bimodal distribution of DNAm patterns in other cell types (Figure S5A).^25^ In addition, not including nRBCs in the reference library, as occurs when using the Gervin reference data set, violates one of the assumptions of the deconvolution method, which is that all major cell types are represented in the reference set.^18^ Thus, we assessed the impact of removing nRBCs from our reference set. For each sample in the validation data set, we predicted cell type proportions with and without the nRBCs included. We then calculated percentage change in estimated proportion for each sample and found an uneven impact across cell types, with B cells (20% mean, 50% maximum), monocyte (10% mean, 52% maximum) and NK (21% mean, 62% maximum) cells having the largest percentage difference in predicted cell type caused by the removal of nRBCs from the reference set (Figure S5A). The magnitude of impact was related to both the abundance of cell type and similarity of DNAm profiles between cord blood cell types as shown by hierarchical clustering across discriminating probes used in the deconvolution (Figure S5B). This demonstrated how inclusion of nRBCs, which displayed distinct DNAm patterns in cord blood, was crucial for accurate deconvolution.

### Using cord references, modified probe selection, and including nRBCs resulted in improved age-specific cell type prediction accuracies in whole blood data sets

Our results have shown that the difference in performance between adult and cord blood were likely not due to any single main factor, but rather the compounding of multiple effects based on the unique properties of cord blood. By resolving the issues identified above, we produced cord blood prediction performance that were more comparable to previously reported adult whole blood deconvolution in our validation data (B cell rho=0.73, CD4T rho=0.84, CD8T rho=0.40, gran rho=0.64, mono rho=0.53, NK rho=0.67, nRBC rho=0.66, Figure 3A).^19^ All cell types showed good correlations, though nRBCs and CD8 T cells seemed to be over-estimated across all samples. PCA analysis shows that most of our predicted cell type proportions were significantly associated with PC1 of cord blood DNAm, as expected (Figure 3B). Interestingly, the signal for nRBCs, specific to cord blood, was only associated with PC2 and PC4, signifying that the nRBC contribution to DNAm pattern in cord blood accounts for less variance than the other cell types.

**Figure 3:**
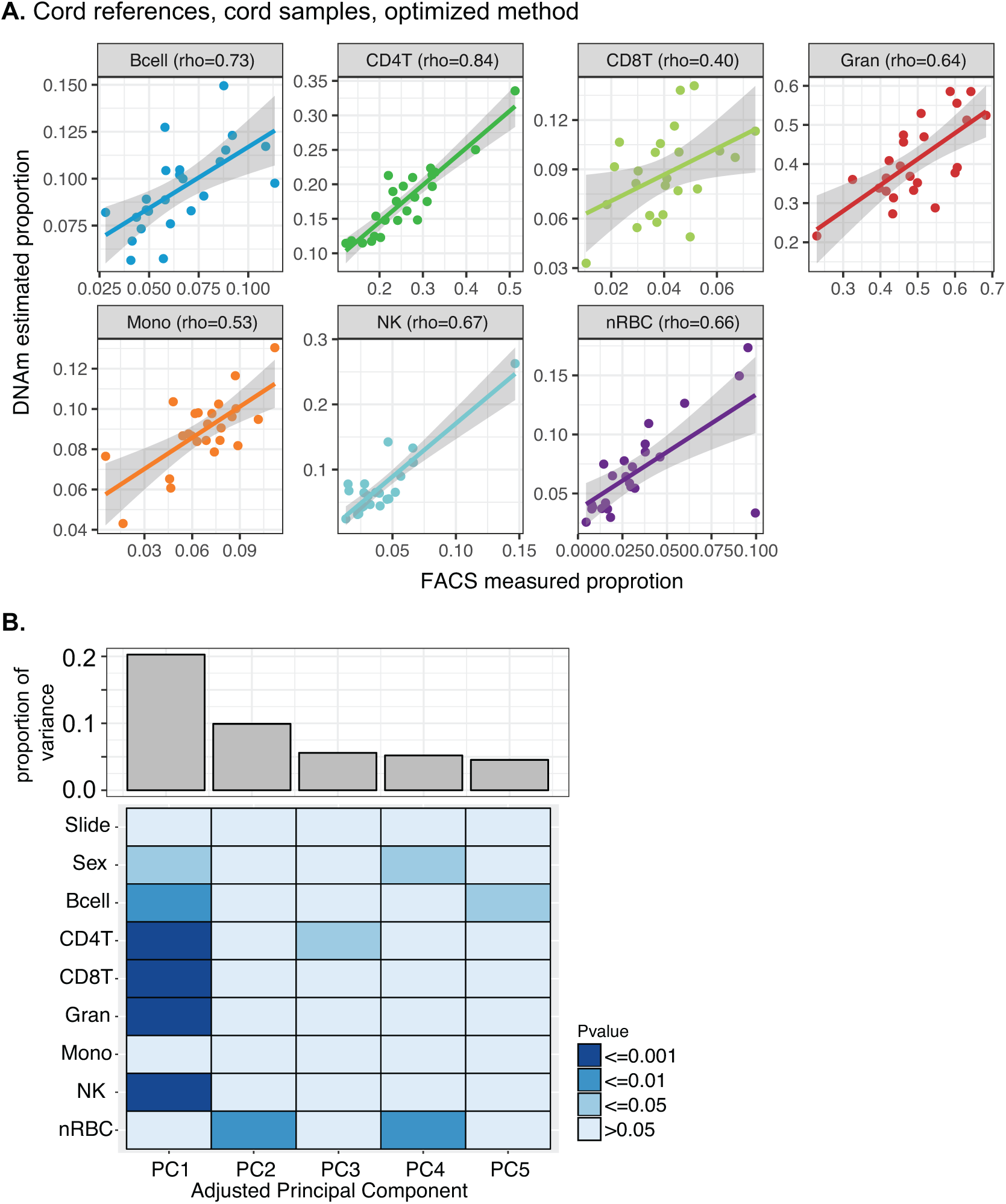
Including nRBCs, altering the probe selection, and using cord references improved deconvolution accuracy in cord blood. A) flow cytometry-based cell counts (x axis) compared with DNAm predicted (y axis) enumeration for each of the seven cord blood cell types. Spearman’s rho for each is shown. Coloured lines indicate regression line for each cell type, and shaded areas 95% confidence interval. B) PCA on 24 whole cord bloods shows associations between deconvolution-based cell counts and PC1, as expected.

To validate the modifications to the cord blood cell type prediction method, we applied our method to an external data set of 191 cord blood samples with flow cytometry-based cell type enumerations from the Generation R cohort study. ^10,20,30^ Unfortunately, this validation data set has not measured nRBCs, and so although we predicted nRBCs, we were unable to validate predictions of this important cell type. However, the prediction accuracy of the other cell types were generally higher than previously shown (Figure 4).^20^ This indicates that the combination of using cord references and including nRBCs combined with correct probe type selection for cord blood result in accurate predictions across data generating platforms.

**Figure 4:**
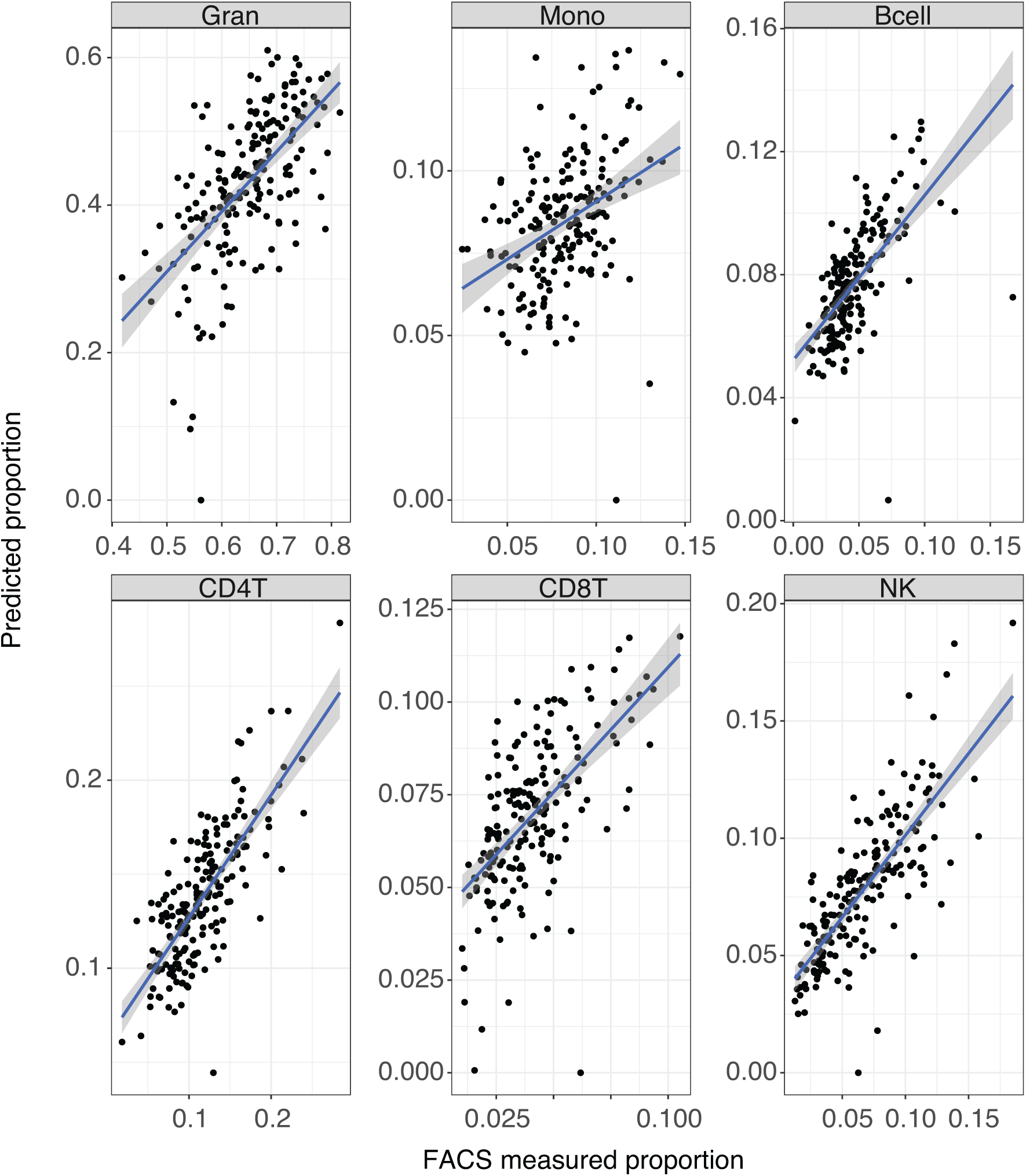
Improved deconvolution resulted in improved prediction accuracy in an independent validation cohort of 191 cord blood samples. Blue line indicates linear regression line, and grey shading indicates 95% confidence interval. nRBCs were predicted using deconvolution but not measured using FACS and so are not shown.

## Discussion

Here, we have explored intrinsic biological differences in DNAm between cord and adult blood cell types. In addition to providing important insights into the fundamental developmental trajectories of DNAm, these analyses led to important adaptations to deconvolution methods that are necessary for accurate predictions of cell type composition in whole cord blood samples. It has been previously shown that DNAm is highly variable with age, but age-related effects on individual cell types have not been as well studied.^2,24^

At least two clear differences exist between cord and adult white blood cells that could influence DNAm, and which might differ across cell types. The first is the impact of age and immunological maturation of white blood cell types after birth.^2,31,32^ While not yet documented, it was perhaps not surprising that DNAm patterns from cord and adult cell types were visually distinct in both clustering and PCA analysis. However, the number of variable sites within cell types was highly different between cord and adult samples. Though all cell types had distinct PCA patterns between cord and adult, this was accentuated in lymphoid than myeloid cells. This observation likely reflected a predominant functional maturation of lymphoid cells post-natally, in contrast to myeloid cells whose function matures predominantly earlier during fetal development. ^10,31,32^ Further, CD4 and CD8 T cells clustered very differently in cord blood than in adult blood. In adults, CD4 and CD8 T cells clustered separately, whereas in cord blood they clustered by individual rather than by cell type. These results suggested that neonatal CD4 and CD8 T cells were more similar at the DNAm level, which may reflect a relative lack of functional differentiation between these two cell lineages prior to antigen exposure. This may also explain why CD8 T cells proportions were difficult to predict accurately in cord blood here, as was also observed in a previous study.^20^ It is possible to predict CD4 T cells more accurately due higher abundance, compared to CD8 T cells, which are both hard to discriminate and of lesser abundance in cord blood.^33^ Thus, a possible solution could be to combine these two cell types for prediction of cell composition in cord blood samples.

Both variability and EWAS results demonstrated that DNAm differences between cord and adult blood were distinct between cell types. Adult cell types were more variable than cord, with B, CD8 T, and NK cells showing the largest differences. EWAS within cell types identified thousands of DNAm differences between cord and adult, but the specific number of differentially methylated CpGs varied across cell types, with granulocytes showing the most differentially methylated sites and monocytes the least. This finding is unusual, given that in the PCA analysis, both granulocytes and monocytes showed more similar broad DNAm patterns between cord and adult than any of the lymphoid cell types. Additionally, there were many more sites that differed between cord and adult and were common between the two myeloid cell types than between the lymphoid cells. This could be a further indication that at least at the level of DNAm, myeloid cell types were more similar to one another than the lymphoid cell types, but it was also possible that the overlap appeared higher because there are only two myeloid cell types, rather than the four lymphoid, and so the overlap may be higher by chance.

Interestingly, in lymphoid cells approximately 10% of the estimated differences between cord and adult overlapped cord blood mQTLs, suggesting a genetic influence on the DNAm variation. In myeloid cells it was much higher, up to 79% in monocytes, suggesting that most of the differences between cord and adult cell types were actually genetic effects. This implies much less DNAm change in monocytes with age compared to other cell types, which is also consistent with previous reports.^34^ Combined with the finding that monocytes, uniquely of all cell types, do not have many probes that are more methylated compared to the other cell types, monocytes seem to be quite different from other cell types in terms of their changes in DNAm with age. This may partly explain why, even after modifying the cord blood prediction, monocytes remain one of the most difficult cell types to predict, with the worst correlation coefficient of any cell type in the GenR data and the second worst in our validation data.

The second major difference between cord and adult blood that might impact deconvolution is the presence of nRBCs. Due to their abundance and unusual DNAm patterns, the presence of nRBCs influences DNAm pattern in cord blood, but to date their impact on estimation of cell type proportions has been poorly understood.^25,35^ We showed that omitting nRBCs from the predictions reduced accuracy of predicting the other cell types. This analysis documented how heavily the constrained projection framework depends on a reference of each major cell type in the mixture. Eliminating one cell type will reduce prediction performance, though it is not always clear which cell type will be allocated to make up for one that is missing, and nRBCs may be particularly prone to this due to their unusual DNAm pattern.^18^ In addition, we note that while the predicted nRBC proportions were well correlated with the measured proportions, these were all scaled proportionally higher. One possible explanation is that the amount of nRBC DNA in a sample is not all contained within nRBCs and thus not reflected in cell counts. As these samples were derived from term births, red blood cells are in the process of extruding their nuclei, potentially leaving acellular DNA in the extracellular material which would then be collected along with the nuclear DNA from intact cells ^36^. Such a process could explain the difference observed, as the deconvolution method is predicting the proportion of nRBC DNA, not cells. Given the high correlation, we are confident that the predictions were accurate for use as corrections, although using the magnitudes of predicted nRBC counts as an outcome should be done with caution. Of additional interest, although nRBCs are common in cord blood, they are not unheard of in adults, with substantial amounts having been reported in anemias, some leukemias, and some cardiac conditions.^37^ In those cases, prediction of cell type composition by DNAm using adult references may demonstrate reduced accuracy, as shown in our experiment removing nRBCs in cord blood.

Putting all of our findings into a bigger perspective, we believe that reference-based prediction techniques are currently the best option for dealing with inter-individual differences in cell type proportions where cell counts are not available. In this case, our findings have shown that biological differences are associated with prediction accuracy, but this approach is not without limitations. First, our reference dataset is based on cell purification using FACS. This technology discriminates based on cell surface markers and does not distinguish between subpopulations within a particular cell type, which may also differ across development.^9^ Second, cell counts for our validation data were quantified using a combination of flow cytometry and the complete blood count. This combination of methods runs the risk of compounding errors from the two methods and thus decreasing accuracy. However, given that we were able to show high accuracy of prediction on samples from two independent data sets, we believe that these counts were sufficiently accurate for correction in EWAS studies. Finally, the adult and cord data came from different data sets and different individuals, which means that both genetic differences and batch effects might inflate our estimate of age-specific differences. To account for genetic differences, we assessed mQTL enrichment of our age-specific findings as described. Batch effects are more difficult, as they are confounded by age, and therefore not possible to specifically remove. However, since there is no expectation that batch effects would be cell-type specific, the comparisons between cell types should be equally affected by batch effects, and thus not bias the interpretation of the findings.

## Conclusions

The exciting potential of epigenetic profiling of cord blood as a marker of *in utero* environmental exposure should be balanced by an understanding of the unique properties of that tissue. Based on our results, it is clear that leukocyte of different lineages mature differently *in utero* and after birth resulting in different DNAm between cord and adult cell types. These appeared to be primarily driven by lymphocytes, which have very similar DNAm profiles in cord blood compared to adults, mirroring their acquisition of immunological memory postnatally upon antigen exposure. These findings suggest an important functional role of DNAm in immune cell maturation during development, and indicate why DNAm-based tools that are generated in adults should be applied to other ages like cord blood with care.

## Methods

### Sample collection

Sorted and validation cord blood samples collected at UBC were collected from term elective caesarian deliveries at BC Women’s Hospital. All mothers gave written informed consent, and protocols were approved by University of British Columbia Children’s & Women’s Research Ethics Board (certificate numbers H07-02681 and H04-70488).

### Purification of cord blood reference panel

Cord blood cell types were purified as previously published.^25^ Briefly, we applied seven whole cord blood samples to Lymphoprep (StemCell Technologies Inc., BC, Canada) density gradient to separate granulocytes from mononuclear cells. Granulocytes were further separated from non-nucleated red blood cells by density gradient. The mononuclear fraction (which include nRBCs) was separated into constituent cell types using a stringent flow cytometry gating strategy, as described previously^25^ on a FACSAriaIII (Becton Dickinson), generating purified populations of monocytes (CD3-, CD19-, CD235-, CD14+), CD4 T cells (CD14-, CD19-, CD235-, CD3+, CD4+), CD8 T cells (CD14-, CD19-, CD235-, CD3+, CD8+), NK cells (CD3-, CD19-, CD235-, CD14-, CD56+), B cells (CD3-, CD14-, CD235-, CD19+), and nucleated red blood cells (CD3-, CD14-, CD19-, CD235+, CD71+).

### Quantification of cell type proportions in whole cord blood validation samples

In addition to the sorted cord blood cell types, we collected twenty-four cord blood samples for validation. A small aliquot of each sample was sent to the BC Children’s Hospital hematology lab for complete blood count with differential (CBC). A second aliquot was prepared as the reference samples above, with the same markers and antibodies, after lysis of red blood cells using BD FACS Lysing Solution (BD Bioscience). Final cell counts for nRBCs, granulocytes, monocytes and lymphocyte subsets were determined combining the CBC and flow cytometry data. CBC provided nRBC, monocyte, and granulocyte cells counts, as well as total lymphocytes. We then scaled these counts to total 1, and calculated lymphocyte subsets (relative proportions of B, NK, CD4T, and CD8T cells) by multiplying the total lymphocyte proportion by the relative proportions of lymphocyte subtypes measured by flow cytometry.

### Generation of DNAm data

DNA from sorted reference and whole validation cord blood samples was isolated using a Qiagen DNAeasy DNA isolation kit (Qiagen, USA). 750ng of isolated DNA was subjected to bisulfite conversion using the Zymo EZDNA bisulfite conversion kit (Zymo Research, USA), then applied to Illumina 450k microarrays per manufacturer’s instructions. Raw data was imported into Illumina Genome Studio (Illumina, USA) for background subtraction and colour correction, then exported into R statistical software for analysis. Reference and validation data were processed and applied to the arrays in separate batches to simulate typical applications.

DNAm preprocessing included removing probes for high detection p-value (> 0.01), low bead coverage (<3), sex chromosomes, cross hybridizing probes, and SNP probes. Next, we applied Noob to correct for background and BMIQ for probe type normalization.^38,39^ Finally, we applied batch correction using ComBat, accounting for chip variability while explicitly protecting cell type.^40,41^ The same protocol was used to normalize the FACS-sorted cord blood data from our study and another study along with the adult sorted data.^3,20,25^ Validation data from Generation R data was generated as previously described.^20,30,42^ For this study, we obtained it as IDAT files and normalized it using the same steps described above.

### Comparison of cord and adult samples

The dendrogram was generated using complete linkage of a Euclidean distance matrix of samples based on methylation beta values including all three of these FACS-sorted cell type data sets, with samples coloured by cell type, age, and myeloid vs lymphoid lineage. Silhouette analysis used the same distance matrix, clustered by cell type. We performed PCA on the same data set using the prcomp function in R. For the number of variable sites, we used only the adult data and our sorted data with similar N, counted the number of sites with SD > 0.05 in beta value in each cell type by each age. To assess pairwise differences between cell types within adult and cord blood, we also used only our sorted cord blood data set with the adult data, and performed a two-group t-test on methylation m-values to determine the number of differentially methylated probes between each pair, using a nominal p value of 1×10^−7^. As DNAm beta values are heteroscedastic, M values are a log transformation of beta values that avoids the typical statistical problems with heteroscedasticity.

We calculated the number of sites which discriminated between cell types and were higher- or lower-methylated in that cell type compared to others by first ranking all sites by p value calculated from a two sided t-test comparing that cell type to all other cell types. Next, we took the top 50 sites that had a mean DNAm value higher in that cell type than others, and the top 50 with a lower mean DNAm value, and plotted the magnitude of mean DNAm difference between that cell type and the other cell types.

### Epigenome-wide association study comparing cord to adult white blood cells

EWAS analysis was performed on the adult and our sorted cord blood data sets.^3,25^ We applied the R package limma with a categorical variable of cord vs adult and no other covariates to normalized data, using a p value cutoff of 1×10^−7^ and a mean absolute beta value difference of 0.1 to define a significant CpGs.

### Cell type prediction

For prediction of cell type proportions in cord blood, we applied the constrained projection quadratic programming (CP/QP) algorithm developed by Houseman et al., Houseman:2012km as implemented in the minfi package without modification, and using either adult or cord reference libraries.^3,5,18,25,29^ We then quantified the sensitivity of our procedure by comparing estimated proportions on the same set of samples depending on whether or not a nRBC profile was available, and after optimized preprocessing and feature selection. Finally, we performed deconvolution again using the cord blood references and defining the sites used in deconvolution as the sites with the top f statistic regardless of direction of change on both our validation and the Generation R data. Accuracy of deconvolution estimates with cell counts was measured with Spearman’s Rho in all cases.

## Declarations

### Ethics approval and consent to participate

The study protocol was approved by the Medical Ethical Committee of the Erasmus Medical Centre, Rotterdam. Written informed consent was obtained for all participants.

### Consent for publication

Not applicable

### Availability of data and material

Reference cord and adult data and used in this study is available on GEO (GSE35069, GSE82084). Validation samples that were internally generated are also available (GEO# TBD), and Generation R data may be available upon request from the study coordinators. Full code for figures is available on GitHub (https://github.com/megjones/Rotterdam_code/blob/master/CordvsAdult_figures), and modified code for cord deconvolution is available https://github.com/megjones/Rotterdam_code/blob/master/Initial_deconvolution_edited. R.

### Authors’ contributions

MJJ conceived the study, performed data analysis, and drafted the manuscript. LD performed data analysis and helped draft the manuscript. HRR collected reference and validation cord blood samples. OdG performed FACS to generate reference samples. JLM and AMM ran the 450k arrays for the de Goede references and internal validation samples. KG, RN, and RL advised on study design and data analysis. LD, MCvZ, and HAM generated FACS data for the GenR cohort. WPR, DCK, JFF, PLM, and SM helped conceive the study, advised on study design, and helped edit the manuscript. MSK helped conceive the study, oversaw the study, and helped edit the manuscript. All authors read and approved the final manuscript.

#### Acknowledgements

The general design of the Generation R Study is made possible by financial support from the Erasmus Medical Center, Rotterdam, the Erasmus University Rotterdam, the Netherlands Organization for Health Research and Development and the Ministry of Health, Welfare and Sport. The Generation R Study is conducted by the Erasmus Medical Center in close collaboration with the School of Law and Faculty of Social Sciences of the Erasmus University Rotterdam, the Municipal Health Service Rotterdam area, Rotterdam, the Rotterdam Homecare Foundation, Rotterdam and the Stichting Trombosedienst & Artsenlaboratorium Rijnmond (STAR-MDC), Rotterdam. We gratefully acknowledge the contribution of children and parents, general practitioners, hospitals, midwives and pharmacies in Rotterdam.

The generation and management of the Illumina 450K methylation array data (EWAS data) for the Generation R Study was executed by the Human Genotyping Facility of the Genetic Laboratory of the Department of Internal Medicine, Erasmus MC, the Netherlands. We thank Mr. Michael Verbiest, Ms. Mila Jhamai, Ms. Sarah Higgins, Mr. Marijn Verkerk and Dr. Lisette Stolk for their help in creating the EWAS database.

## Supplementary figures and tables

**Figure S1:**
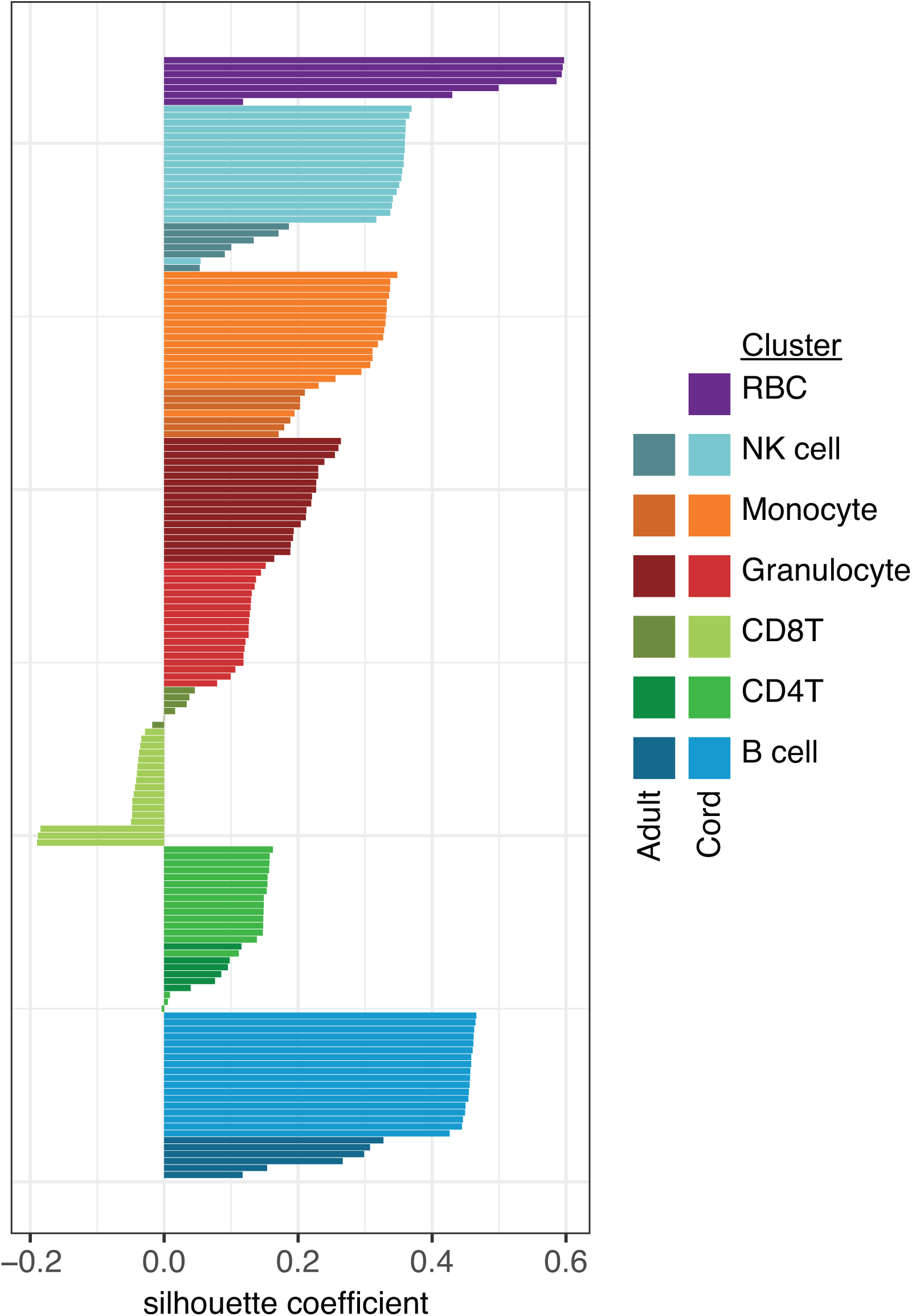
DNA methylation patterns of leukocyte cell types cluster together. Silhouette plot based on DNAm data show 7 clusters. Cord and adult samples are indicated in the same colour, but different shades (dark for adult and light for cord blood data).

**Figure S2:**
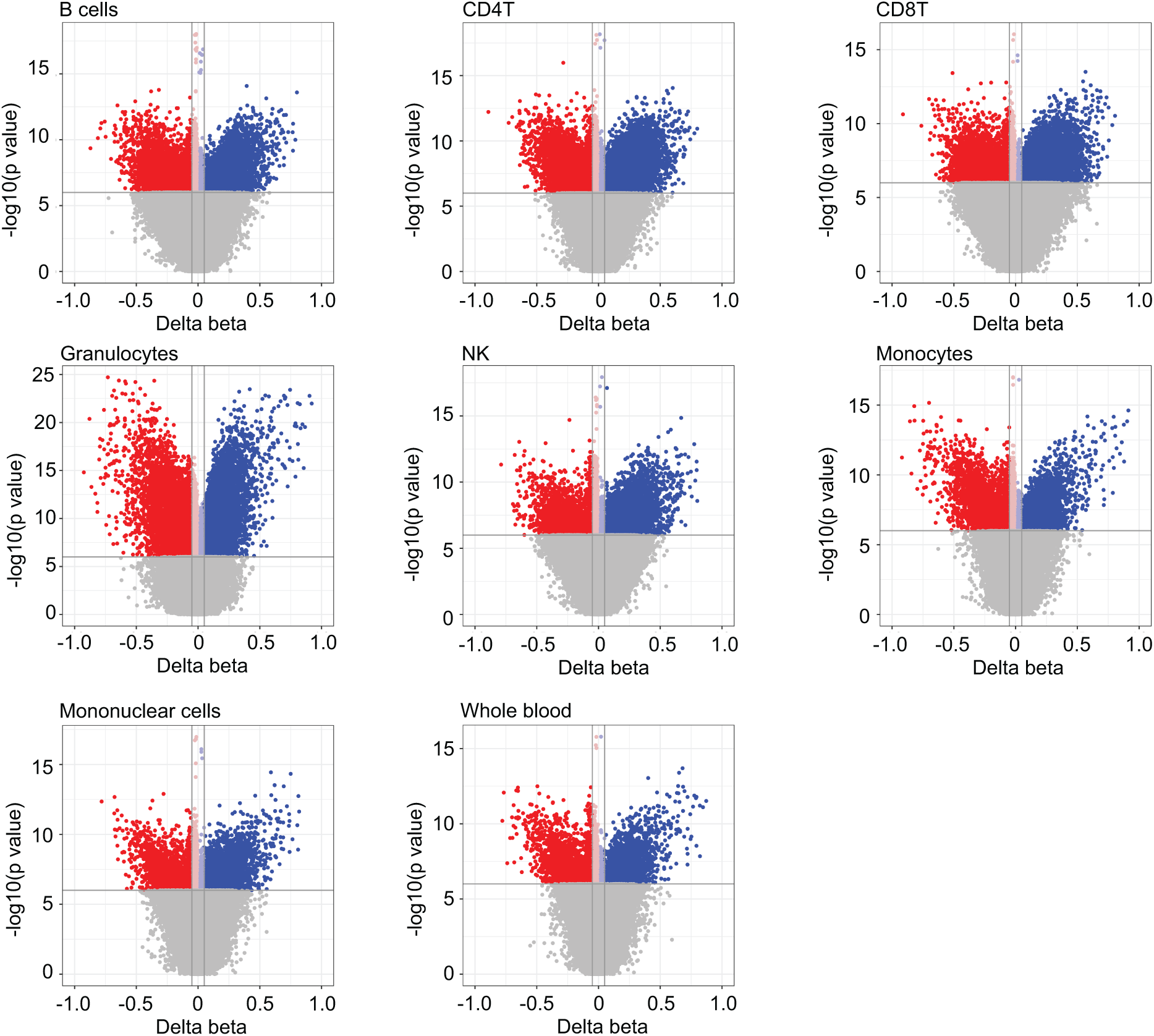
EWAS analysis shows large differences between cord and adult in all cell types. Volcano plots for each component cell type and two cell mixtures (whole blood and mononuclear cells). Sites in grey did not meet the 1×10^−7^ p value cutoff. Sites in light red and light blue did not meet the absolute beta value difference of 0.1.

**Figure S3:**
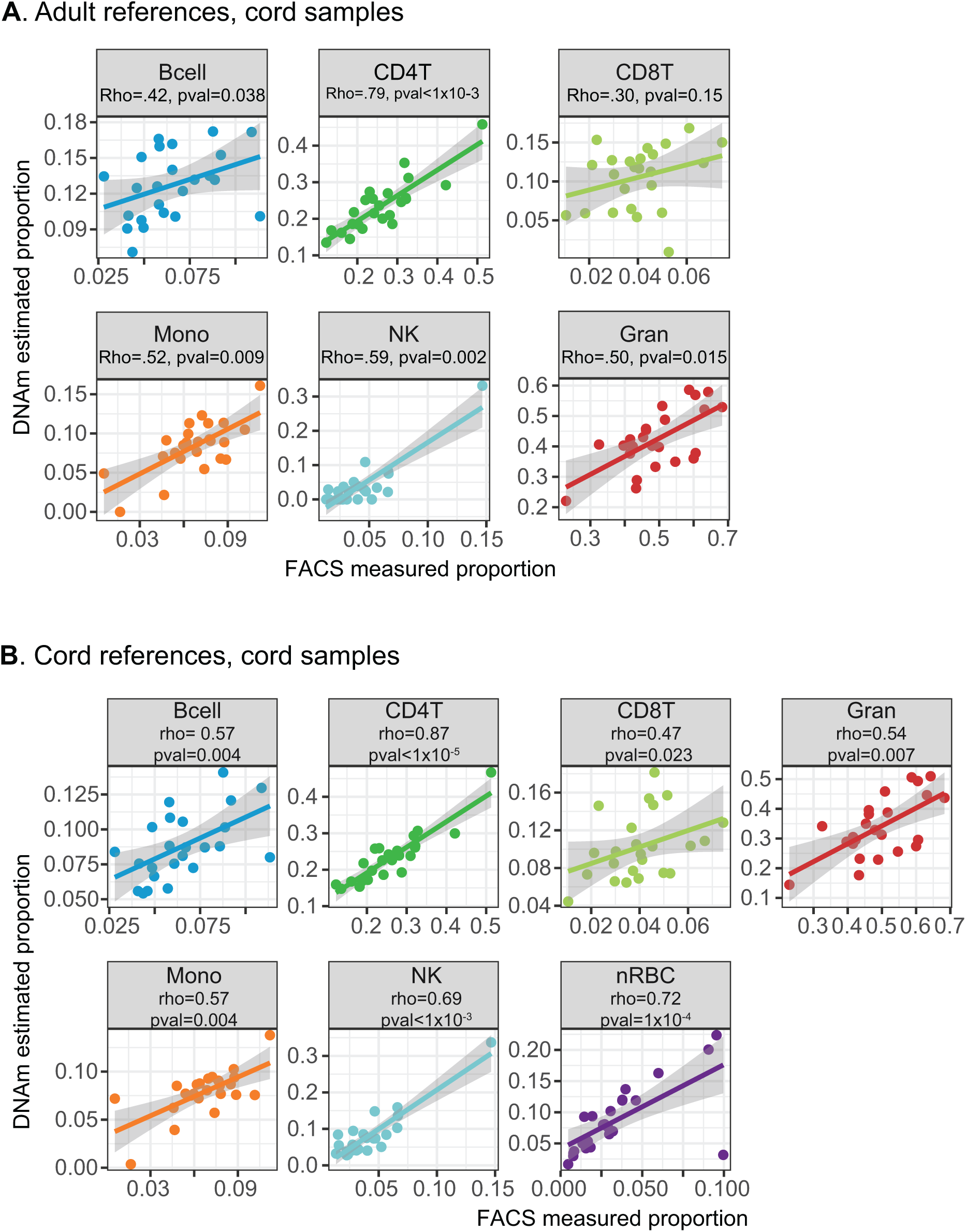
Using adult references in deconvolution of cord blood results in poor prediction, and using cord references improves predictions, but some cell types remain poorly predicted. In 24 cord blood cell types, flow cytometry-based cell counts (x axis) are plotted against DNAm deconvolution-based estimates (y axis), using either adult **(A)** or cord **(B)** blood references.

**Figure S4:**
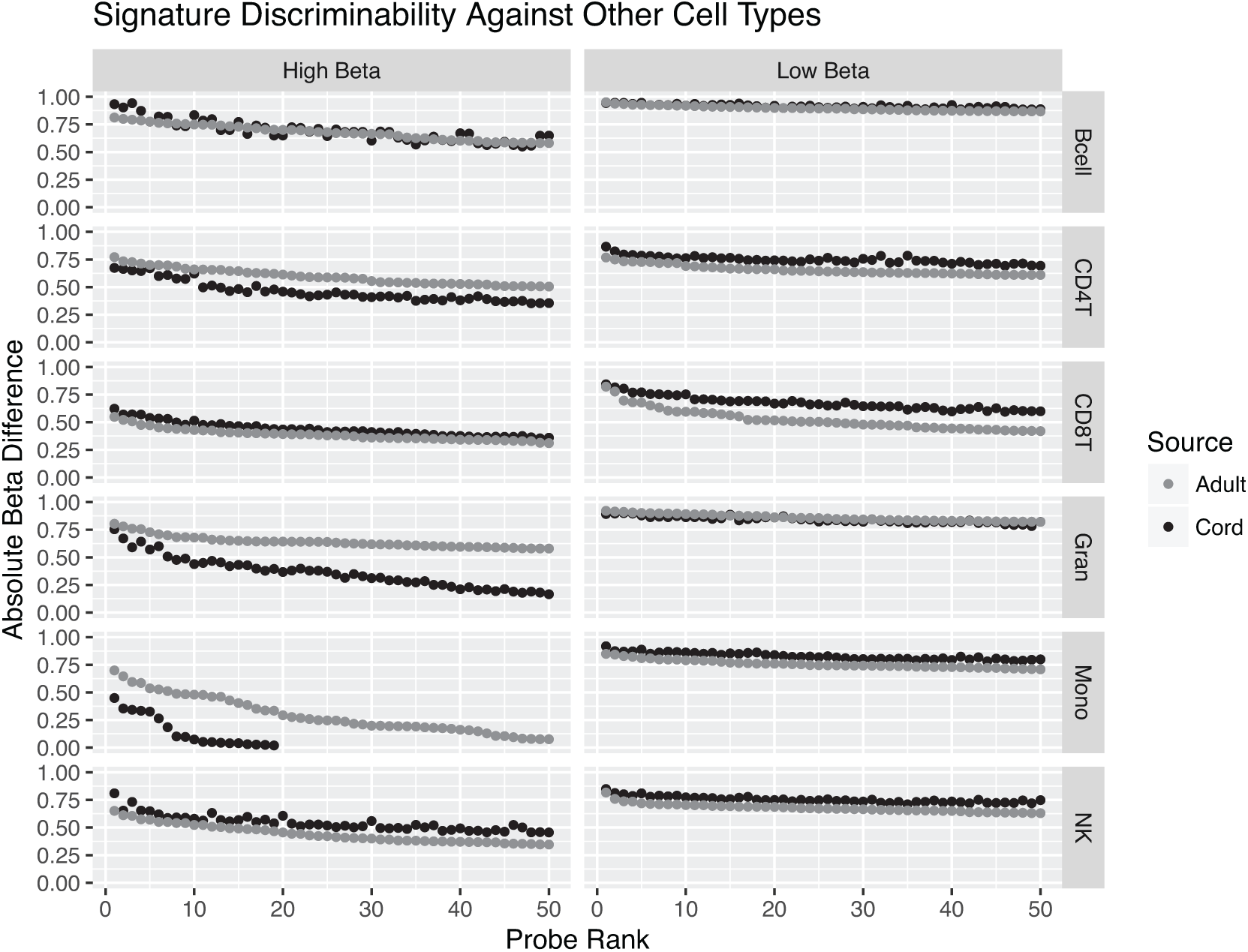
Cord blood had fewer cell type-distinguishing CpGs that were more methylated in a particular cell type than other cell types. Plots show the 50 best ranked distinguishing probes by p value (x axis) versus the average DNAm difference between a particular cell type and other cell types (y axis), and whether they are more (top) or less (bottom) methylated in that cell type than others. For each cell type, the sites that are more methylated drop to 0 in actual DNAm difference before reaching 50, meaning that some of the probes that would have been chosen to use in deconvolution are actually not differentially methylated in that cell type at all.

**Figure S5:**
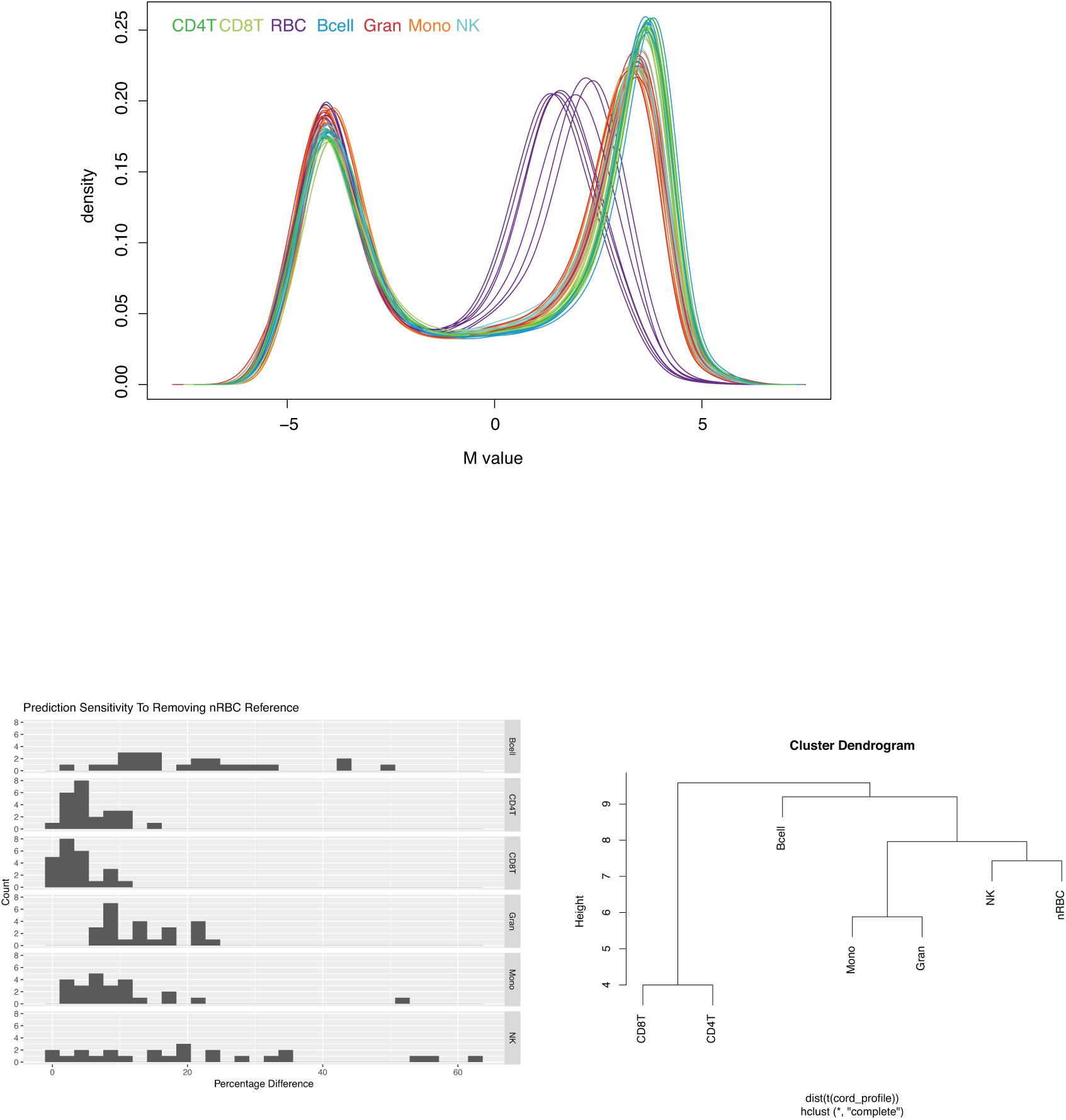
Including nRBCs in deconvolution of cord blood is important for accurate predictions of all cell types. **A)** DNAm profiles for cord white blood cell types. N=7 for each cell type except CD8 T cells, where N=6. **B)** Histogram showing the difference between predicted and actual cell counts for each cell type if nRBCs were not included in the prediction. **C)** Dendrograms showing relationships in DNAm pattern at sites used in deconvolution across the 7 cord blood cell types. NK cells are the most similar to nRBCs at these sites, explaining why this cell type is the most impacted by not including nRBCs.

**Table S1:**
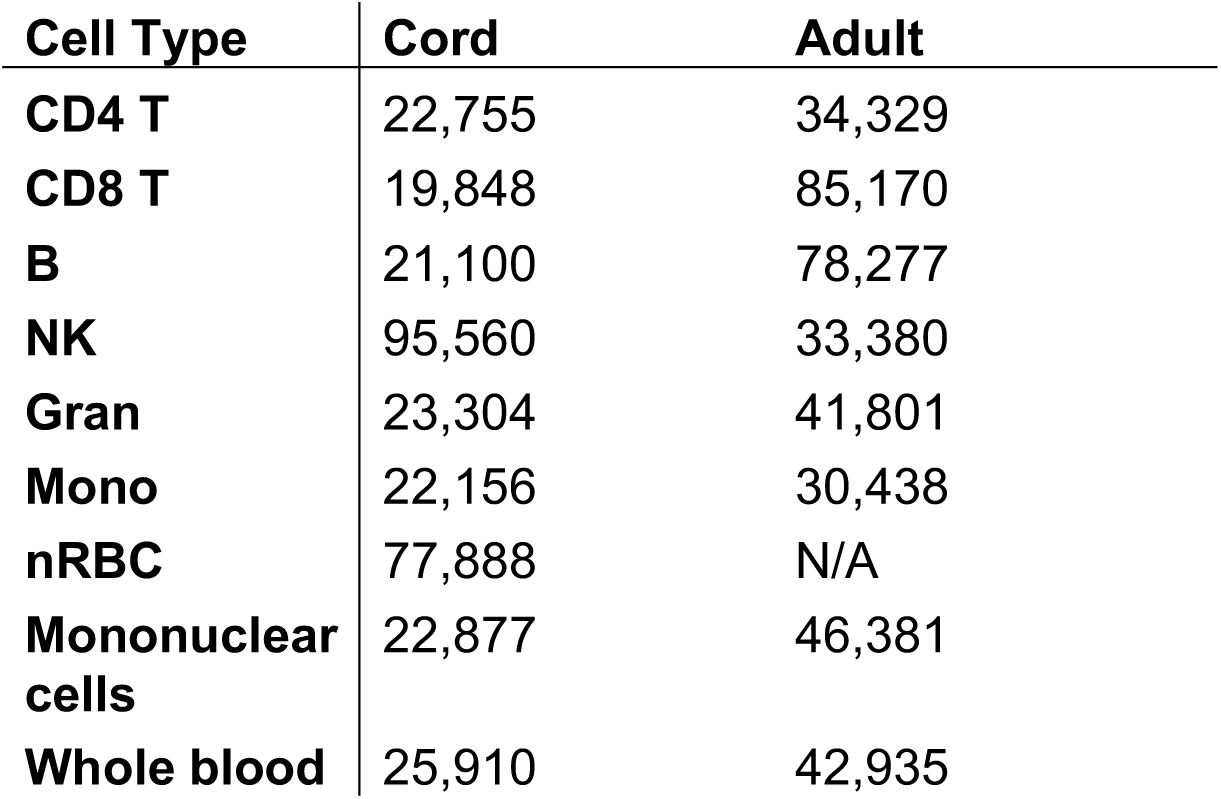
Numbers of variable CpGs, defined as SD>0.05 in cord and adult white blood cell types

**Table S2:** Number of differentially methylated sites between cord and adult and how many of those are mQTLs

|  | **N unique DM sites** | **N mQTLs** | **Percentage mQTLs** |
| --- | --- | --- | --- |
| **B** | 6393 | 387 | 6.05 |
| **CD4 T** | 7840 | 636 | 8.11 |
| **CD8 T** | 4443 | 377 | 8.48 |
| **G** | 7235 | 1355 | 18.72 |
| **Mo** | 339 | 268 | 79.05 |
| **NK** | 3518 | 344 | 9.78 |
| **All cell types** | 588 | 65 | 11.05 |
| **Myeloid** | 2062 | 242 | 11.73 |
| **Lymphoid** | 397 | 37 | 9.31 |

**Table S3:** Pairwise differentially methylated cord vs adult probe overlaps

|  | **B** | **CD4T** | **CD8T** | **G** | **Mo** | **NK** |
| --- | --- | --- | --- | --- | --- | --- |
| **B** | **7378** |  |  |  |  |  |
| **CD4 T** | 2344 | **8825** |  |  |  |  |
| **CD8 T** | 2079 | 3071 | **5428** |  |  |  |
| **G** | 2558 | 2595 | 1746 | **9885** |  |  |
| **Mo** | 1472 | 1342 | 970 | 2650 | **2989** |  |
| **NK** | 2202 | 2140 | 1987 | 2081 | 1166 | **4503** |

